# Improved vectors and genome-wide libraries for CRISPR screening

**DOI:** 10.1101/006726

**Authors:** Neville E. Sanjana, Ophir Shalem, Feng Zhang

## Abstract

Genome-wide, targeted loss-of-function pooled screens using the CRISPR (clustered regularly interspaced short palindrome repeats)–associated nuclease Cas9 in human and mouse cells provide an alternative screening system to RNA interference (RNAi) and have been used to reveal new mechanisms in diverse biological models^1–4^. Previously, we used a Genome-scale CRISPR Knock-Out (GeCKO) library to identify loss-of-function mutations conferring vemurafenib resistance in a melanoma model^1^. However, initial lentiviral delivery systems for CRISPR screening had low viral titer or required a cell line already expressing Cas9, limiting the range of biological systems amenable to screening.

---

Here, we sought to improve both the lentiviral packaging and choice of guide sequences in our original GeCKO library^1^, where a pooled library of synthesized oligonucleotides was cloned into a lentiviral backbone containing both the *Streptococcus pyogenes* Cas9 nuclease and the single guide RNA (sgRNA) scaffold. To create a new vector capable of producing higher-titer virus (lentiCRISPRv2), we made several modifications, including removal of one of the nuclear localization signals (NLS), human codon-optimization of the remaining NLS and P2A bicistronic linker sequences, and repositioning of the U6-driven sgRNA cassette (**Fig. 1a**). These changes resulted in a ∼10-fold increase in functional viral titer over lentiCRISPRv1^1^ (**Fig. 1b**).

**Figure 1.**
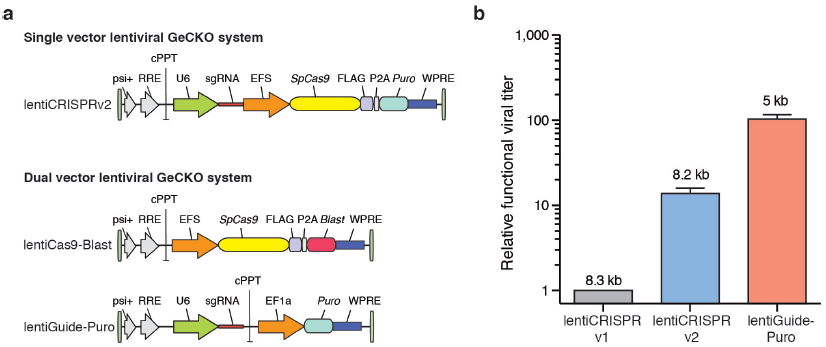
New lentiviral CRISPR designs produce viruses with higher functional titer. (**a**) Lentiviral expression vector for *Streptococcus pyogenes* Cas9 and sgRNA in the improved one vector system (lentiCRISPR v2) and the two vector system (lentiCas9-Blast, lentiGuide-Puro). Psi packaging signal (psi+), rev response element (RRE), central polypurine tract (cPPT), elongation factor-1α short promoter (EFS), FLAG octapeptide tag (FLAG), 2A self-cleaving peptide (P2A), puromycin selection marker (puro), posttranscriptional regulatory element (WPRE), blasticidin selection marker (blast), and elongation factor-1α promoter (EF1a). (**b**) Relative functional viral titer of viruses made from lentiCRISPR v1^1^, lentiCRISPR v2, and lentiGuide-Puro vector with a EGFP-targeting sgRNA (*n* = 3 independently-transfected virus batches with 3 replicate transductions per construct). HEK293FT cells were transduced with serial dilutions of virus and, after 24 hours, selected using puromycin (1 ug/ml). Puromycin-resistant cells were measured after 4 days from the start of selection using the CellTiter-Glo (Promega) luciferase assay. Relative titers were calculated using viral volumes that yielded less than 20 % puromycin-resistant cells in order to minimize the number of cells with multiple infections. Numbers above each bar indicate the size of the packaged virus for each construct.

To further increase viral titer, we also cloned a two-vector system, in which Cas9 (lentiCas9-Blast) and sgRNA (lentiGuide-Puro) are delivered using separate viral vectors with distinct antibiotic selection markers (**Fig. 1a**). LentiGuide-Puro has a ∼100-fold increase in functional viral titer over the original lentiCRISPRv1 (**Fig. 1b**). Both single and dual-vector systems mediate efficient knock-out of a genomically-integrated copy of EGFP in human cells (**Supplementary Fig. 1**). Whereas the dual vector system enables generation of Cas9-expressing cell lines which can be subsequently used for screens using lentiGuide-Puro, the single vector lentiCRISPRv2 may be better suited for *in vivo* or primary cell screening applications.

**Supplementary Figure 1.**
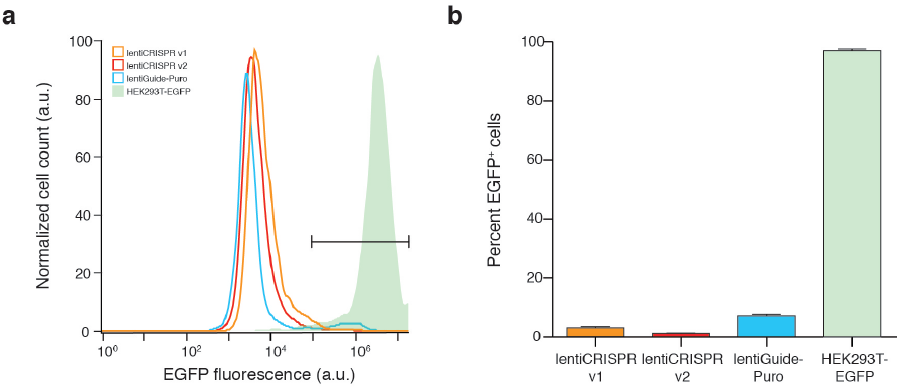
EGFP expression in HEK293T-EGFP cells after transduction with different lentiviral CRISPR constructs. (**a**) Representative histograms of EGFP fluorescence from single transductions of HEK293T-EGFP cells with lentiCRISPR v1, lentiCRISPR v2, lentiGuide-Puro or no virus. For lentiGuide-Puro, HEK293T-EGFP cells had previously been transduced with lentiCas9-Blast and selected for 4 days with blasticidin. Twenty-four hours after transduction, cells were selected in puromycin and then analyzed by flow cytometry at 7 days after infection. (**b**) Percentage of EGFP positive cells (as given by gate drawn in **a**) per viral construct (error bars indicate s.e.m, *n* = 3 biological replicate transductions).

In addition to the vector improvements, we designed and synthesized new human and mouse GeCKOv2 sgRNA libraries (**Supplementary Methods**) with several improvements (**Table 1**): First, for both human and mouse libraries, to target all genes with a uniform number of sgRNAs, we selected 6 sgRNAs per gene distributed over 3-4 constitutively expressed exons. Second, to further minimize off-target genome modification, we improved the calculation of off-target scores based on specificity analysis^5^. Third, to inactivate microRNAs (miRNAs) which play a key role in transcriptional regulation, we added sgRNAs to direct mutations to the pre-miRNA hairpin structure^6^. Finally, we targeted ∼1000 additional genes not included in the original GeCKO library.

**Table 1.** Comparison of new GeCKO v2 human and mouse sgRNA libraries with existing CRISPR libraries.

|  | Wang et al. (2014)<br>library | Shalem et al. (2014)<br>GeCKO v1 library | Yusa et al. (2014)<br>library | GeCKO v2<br>human library | GeCKO v2<br>mouse library |
| --- | --- | --- | --- | --- | --- |
| *Species* | human | human | mouse | human | mouse |
| *Number of genes targeted* | 7,114 | 18,080 | 19,150 | 19,050 | 20,611 |
| *Targeting constructs per gene* | 10 per gene | variable (typically 3 to 4 per gene) | variable (typically 4 to 5 per gene) | 6 per gene<br>(3 in Library A,<br>3 in Library B) | 6 per gene<br>(3 in Library A,<br>3 in Library B) |
| *Number of miRNA targeted* | none | none | none | 1,864 | 1,175 |
| *Targeting constructs per miRNA* | n/a | n/a | n/a | 4 per miRNA | 4 per miRNA |
| *Control (non-targeting) sgRNAs* | 100 | none | none | 1,000 | 1,000 |
| *Total sgRNA constructs* | 73,151 | 64,751 | 87,897 | 123,411<br>(65,383 in Library A,<br>58,028 in Library B) | 130,209<br>(67,405 in Library A,<br>62,804 in Library B) |
| *Viral plasmid vector* | Dual vector:<br>sgRNA only | Single vector:<br>Cas9 and sgRNA<br>(lentiCRISPR v1) | Dual vector:<br>sgRNA only | Single and dual vector:<br>lentiCRISPR v2 and<br>lentiGuide-Puro | Single and dual vector:<br>lentiCRISPR v2 and<br>lentiGuide-Puro |

New GeCKO libraries target an increased number of genes with uniform coverage (6 sgRNAs per gene divided into sub-libraries with 3 sgRNAs per gene). The GeCKO v2 libraries also contain sgRNAs that target microRNAs and control sgRNAs that do not target in the genome. The libraries have been cloned into lentiCRISPR v2 (which includes Cas9 in the same vector) and lentiGuide-Puro (sgRNA-only vector) and are available in either format.

Both libraries, mouse and human, are divided into 2 sub-libraries — containing 3 sgRNAs targeting each gene in the genome, as well as 1000 non-targeting control sgRNAs. Screens can be performed by combining both sub-libraries, yielding 6 sgRNAs per gene, for higher coverage. Alternatively, individual sub-libraries can be used in situations where cell numbers are limiting (eg. primary cells, *in vivo* screens). The human and mouse libraries have been cloned into lentiCRISPRv2 and into lentiGuide-Puro and deep sequenced to ensure uniform representation (**Supplementary Fig. 2**, **3**). These new lentiCRISPR vectors and human and mouse libraries further improve the GeCKO reagents for diverse screening applications. Reagents are available to the academic community through Addgene and associated protocols, support forums, and computational tools are available via the Zhang lab website (www.genome-engineering.org).

**Supplementary Figure 2.**
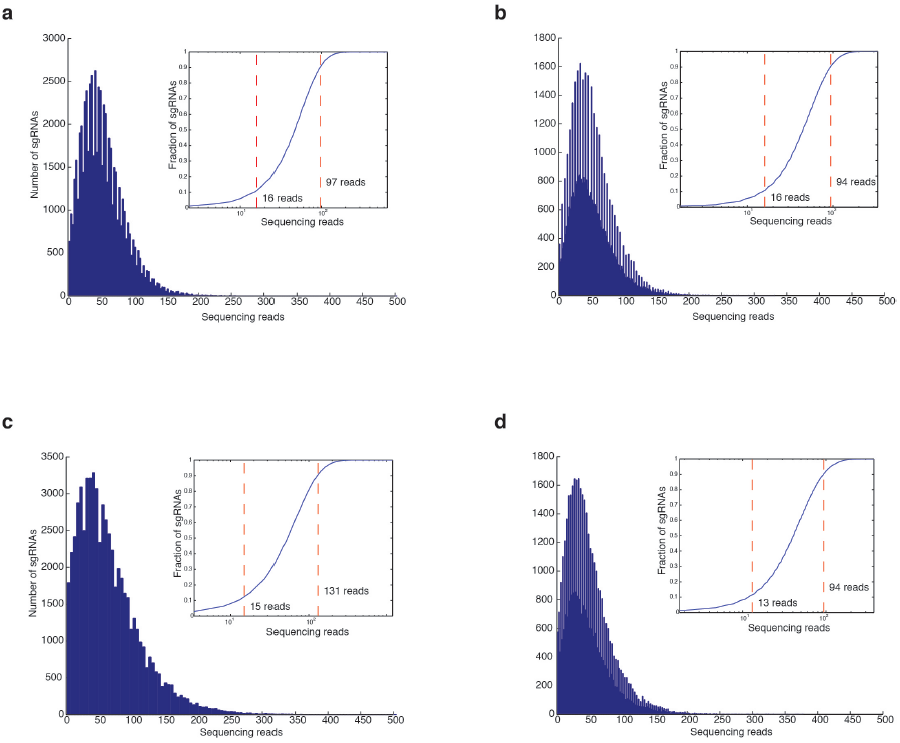
Sequencing of human GeCKOv2 libraries cloned into lentiCRISPRv2 and lentiGuide-Puro. Histograms of sgRNA representation after cloning each half-library into sgRNA vectors. Inset: Cumulative distribution of sequencing reads. The number of sequencing reads for the 10^th^ and 90^th^ sgRNA percentile is indicated by the dashed red lines and text labels. All library sequencing was performed with PCR replicates; only one replicate is shown for each half-library. (**a,b**) Human GeCKOv2 library A (**a**) and library B (**b**) in lentiCRISPRv2. (**c,d**) Human GeCKOv2 library A (**c**) and B (**d**) in lentiGuide-Puro.

**Supplementary Figure 3.**
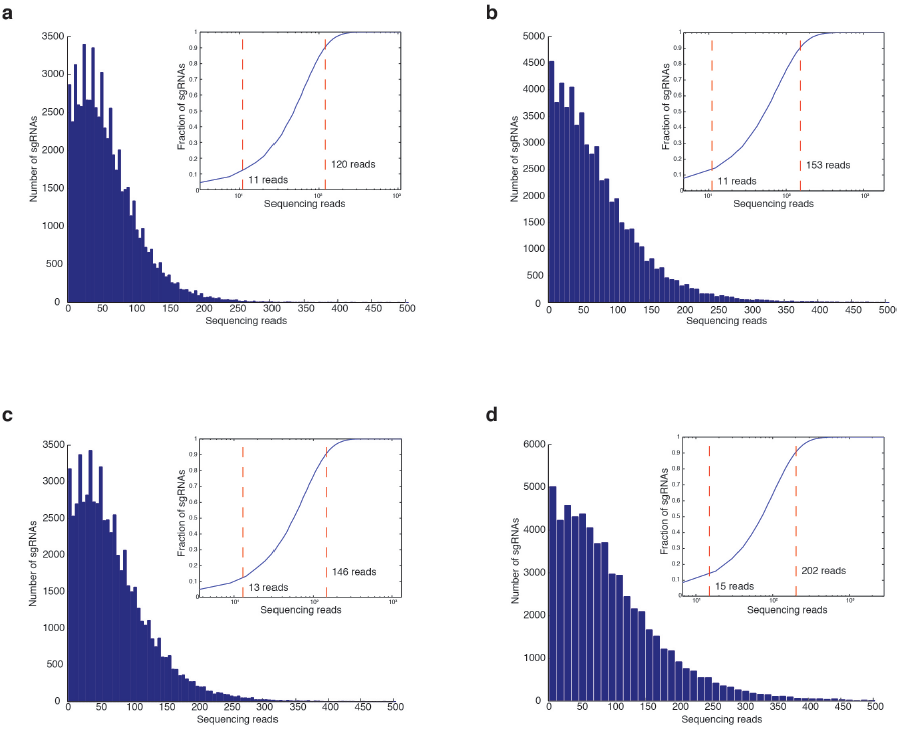
Sequencing of mouse GeCKOv2 libraries cloned into lentiCRISPRv2 and lentiGuide-Puro. Histograms of sgRNA representation after cloning each half-library into sgRNA vectors. Inset: Cumulative distribution of sequencing reads. The number of sequencing reads for the 10^th^ and 90^th^ sgRNA percentile is indicated by the dashed red lines and text labels. All library sequencing was performed with PCR replicates; only one replicate is shown for each half-library. (**a,b**) Mouse GeCKOv2 library A (**a**) and library B (**b**) in lentiCRISPRv2. (**c,d**) Mouse GeCKOv2 library A (**c**) and B (**d**) in lentiGuide-Puro.

## Acknowledgements

We also would like to thank the entire Zhang Lab for support and helpful advice. N.S. is supported by a postdoctoral fellowship from the Simons Center for the Social Brain at MIT. O.S. is a Klarman Fellow of the Broad-Israel Partnership. F.Z. is supported by the NIH and NIMH through a Director’s Pioneer Award (5DP1-MH100706), a Transformative R01 grant (5R01-DK097768), the Keck, Merkin, Vallee, Damon Runyon, Searle Scholars, Klingenstein, and Simons Foundations, Bob Metcalfe, and Jane Pauley.

## Author Contributions

N.S., O.S., and F.Z. conceived and designed the experiments. N.S. and O.S. performed the experiments and analyzed the data. N.S., O.S., and F.Z. wrote the manuscript.

## Supplementary Methods

### Lentiviral cloning and production

For determination of lentiCRISPR v1, lentiCRISPR v2, and lentiGuide-Puro viral titers, the following sgRNA targeting EGFP (with no known targets in the human genome) was cloned into all 3 lentiviral transfer vectors:

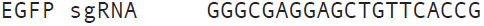

To clone the sgRNA guide sequence, plasmids were cut and dephosporylated with FastDigest BsmBI and FastAP (Fermentas) at 37 °C for 2 hours. Oligonucleotides for the EGFP sgRNA guide sequence (Integrated DNA Technologies) were phosphorylated using polynucleotide kinase (Fermentas) at 37 °C for 30 minutes and then annealed by heating to 95 °C for 5 minutes and cooling to 25 °C at 1.5 °C/minute. Using T7 ligase (Enzymatics), annealed oligos were ligated into gel purified vectors (Qiagen) at 25 °C for 5 minutes. Cloned transfer plasmids were amplified using a endotoxin-free midi-prep kit (Qiagen).

To make lentivirus, the transfer plasmids were co-transfected with packaging plasmids pMD2.G and psPAX2 (Addgene plasmids 12259 and 12260), as described previously^1^. Briefly, for each virus, a T-75 flask of 80 % confluent HEK293T cells (ATCC CRL-3216) was transfected in OptiMEM (Life Technologies) using 10 ug of the transfer plasmid, 5 ug pMD2.G, 7.5 ug psPAX2, 100 ul of Plus Reagent (Life Technologies), and 50 ul of Lipofectamine 2000 (Life Technologies). After 6 hours, media was changed to D10 media, DMEM (Life Technologies) with 10 % fetal bovine serum (Hyclone), with 1 % bovine serum albumin (Sigma) added to improve virus stability. After 60 hours, viral supernatants were harvested and centrifuged at 3,000 rpm at 4 °C for 10 min to pellet cell debris. The supernatant was filtered through a 0.45 um low protein binding membrane (Millipore) and used immediately.

### Lentiviral functional titration

Lentiviruses were titered in a functional assay by measuring puromycin resistance after transduction. All lentiviruses were produced in triplicate transfections. Relative titer measurements from separate viral production transfections performed weeks apart resulted in similar relative titers.

For each viral construct, 2.5×10^4^ HEK293T-EGFP cells were transduced in suspension (i.e. during plating) with 10, 100, or 1000 ul of viral supernatant in wells of a 24-well plate. For lentiGuide-Puro transduction the HEK293T-EGFP cells also had a genomically-integrated copy of Cas9 from previous transduction with lentiCas9-Blast. Each transduction condition (construct and virus volume) was performed in triplicate. In each well, D10 culture media was added to make the final volume 1.5 ml. Cell without any virus added were also plated in six wells (3 wells for puromycin treatment, 3 wells as control).

At 24 hours post-transduction, media was changed to D10 with 1 ug/ml puromycin (Sigma) for all wells except the uninfected controls without puromycin. At 3 days post-transduction, cells in all wells were split 1:5 to prevent any well from reaching confluence. Except for the uninfected controls without puromycin, new D10 media was supplemented with 1 ug/ul puromycin. At 5 days post-transduction, all cells in the uninfected control wells treated with puromycin were floating/dead, which was verified using Trypan Blue exclusion (Sigma).

For the remaining wells, adherent cells were present and cell viability was measured using CellTiter Glo (Promega) following the manufacturer’s protocol. After allowing cells to reach room temperature, media was aspirated from the cells and CellTiter Glo (diluted 1:1 in phosphate-buffered saline) was added. Plates were covered with foil, placed on an orbital shaker for 2 min, and then incubated for 10 minutes at room temperature. Luminescence was measured on an Synergy H4 plate imager (Biotek) using a 1 second integration time and auto-gain to utilize the full dynamic range of the detector. Positive controls (untranduced cells without puromycin) and negative controls (empty wells) were included in the assay.

Fold differences in titer between viral constructs were calculated using luminescence values. Specifically, comparisons were made between pairs of viruses for the same volume of supernatant. Only viral volumes for which cell survival was greater than 1 % and less than 20 % of control (untransduced) cells were directly compared. Assuming Poisson statistics, 20 % cell survival implies that approximately 90 % of cells surviving puromycin selection were infected by only a single virus.

Flow cytometry data was collected from the same set of infections using a BD Accuri C6 flow cytometer. Using FlowJo (TreeStar), single cells were distinguished from debris and doublets by gating in forward and side scatter area plots. EGFP fluorescence was measured in the gated population from transduced and uninfected HEK293T-EGFP cells. For transductions with lentiGuide-Puro, HEK293T-EGFP cells had been transduced with lentiCas9-Blast and selected for 6 days with 5 ug/ml blasticidin (Life Technologies). For all antibiotic selections (puromycin and blasticidin), uninfected control cells were also exposed to the same concentration and duration of antibiotic selection to verify complete elimination of untransduced cells.

### Design of GeCKOv2 libraries

Genome-wide sgRNA libraries for the human and mouse genomes were designed using the following steps:

1. *Identification of constitutive exons:* For the human library, RNA sequencing data from the Illumina Human Body Map 2.0 (GEO accession number: GSE30611) was mapped to the reference human genome (hg19) using TopHat v1.0.14^2^ and transcripts were reconstructed with Cufflinks v1.0.2^3^, as previously described^4^. Exons expressed in all tissues in the Human Body Map dataset were termed constitutive and chosen for sgRNA targeting. For the mouse library, we selected exons that are included in all of the NCBI RefSeq (http://www.ncbi.nlm.nih.gov/refseq/) transcripts for the same gene. In both species, for each gene, we excluded the first and last exons and any exon that contained an alternative splicing site. Using these procedures, we chose 4 constitutive exons for each gene. For genes where there were less than 4 constitutive exons, we added exons starting from the second coding exon towards the end of the gene. The coding boundaries for each exon were identified using the NCBI Consensus CoDing Sequence (CCDS) database (http://www.ncbi.nlm.nih.gov/CCDS/CcdsBrowse.cgi).
2. *Choice of sgRNA sequences:* For each gene, all *S. pyogenes* Cas9 sgRNA sequences of the form (N)_20_NGG in all constitutive exons (from the procedure in (1) above) were selected as candidate targets. Each 20mer candidate target sgRNA was mapped to a precompiled index containing all 20mer sequences in the human genome followed by either NGG or NAG. This mapping was done using the Bowtie short read aligner^5^, allowing up to 3 base mismatches (with parameters -a --best -v 3). For each potential off-target identified by the mapping, the following score was calculated^6^:

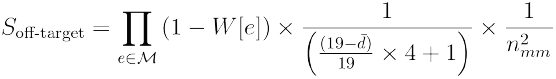 Within the first term, *M* is the set of mismatches between the potential off-target and the candidate target sgRNA. For each mismatch in *M*, *e* denotes the position of the mismatch relative to the protospacer-adjacent motif (PAM). The values of *e* range between 1 (for the most PAM-distal base in the sgRNA) to 20 (for the most PAM-proximal base in the sgRNA). Using *e* as the index, a value from the look-up table *W* denotes the empirically-determined weight (relative to the other terms in the *S*_off-target_ metric) that penalizes mismatches based on their position in the sgRNA sequence^6^:

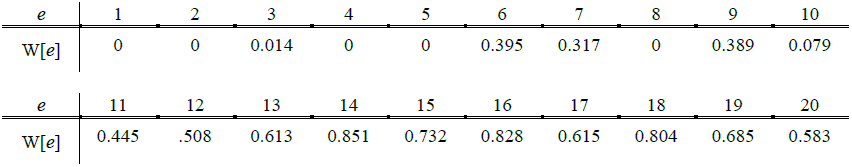 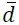 is the mean pairwise distance between all pairs of elements in *M* and *n*_*mm*_ is the number of elements in *M*. Each of the individual off-targets (*S*_off-target_) for one candidate target sgRNA were integrated into an aggregate off-target score as follows:

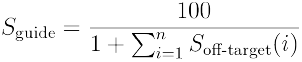 *S*_off-target_ is the score for a single potential off-target and *S*_guide_ is a score (0-100 scale) that provides a rating of how well a candidate sgRNA uniquely targets within the genome. Candidate sgRNAs with higher values of *S*_guide_ are predicted to have less off-target activity. For each gene, we chose 6 sgRNAs with the highest values of *S*_guide_ across all 4 constitutive exons (from step (1)) subject to constraint that no more than 2 sgRNAs could target a single exon. This procedure resulted in extremely uniform coverage of genes: For 99.4 % of genes in the GeCKOv2 human library and 99.8 % of genes in the GeCKOv2 mouse library, each gene was targeted with exactly 6 sgRNAs.
3. *Targeting of mature miRNAs:* For the design of miRNA targeting guides, we used the pre-miRNA sequence coordinates from the mirBASE database (www.mirbase.org)^7^. For each pre-miRNA, we listed all the possible (N)_20_NGG sequences. In total, we chose up to 4 sgRNAs per miRNA or less if there were not 4 sgRNAs present in the pre-miRNA sequence.

### GeCKO library pooled synthesis and cloning

DNA oligonucleotide library synthesis was completed on a programmable microarray using a B3 Synthesizer (CustomArray) and SAFC Proligo reagents (Sigma), as recommended by the manufacturer. The synthesis products were cleaved from the microarray and deprotected by overnight incubation in 28-30 % ammonium hydroxide at 65 °C, dried, resuspended in 30 ul TE buffer and then purified using a QIAquick spin column (Qiagen). Full-length oligonucleotides (74 nt) were amplified by PCR using Phusion HS Flex (NEB). For the PCR reaction, the manufacturer’s protocol was followed using 0.1 ul of synthesized oligonucleotide template, primers Array F and ArrayR (see below), an annealing temperature of 63 °C, an extension time of 15 s, and 20 cycles. After PCR, the 140 bp amplicon was size-selected using a 2 % agarose E-Gel EX (Life Technologies, Qiagen).

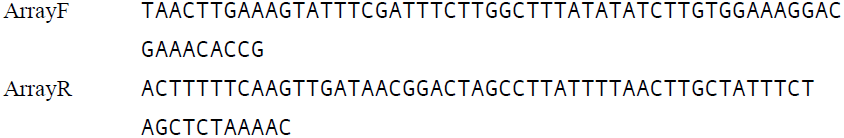

The vector backbone (lentiCRISPR v2 or lentiGuide-Puro) was digested with BsmBI (Fermentas) and treated with FastAP (Fermentas) at 37 °C for 2 hours and gel-purified on a 1 % E-Gel EX (Life Technologies, Qiagen). A 20 ul Gibson ligation reaction (NEB) was performed using 10 ng of the gel-purified inserts and 25 ng of the vector (for lentiCRISPR v2) and using 5 ng of the gel-purified inserts and 25 ng of the vector (for lentiGuide-Puro). From the ligation, 0.5 ul of the reaction was transformed into 25 ul of electrocompetent cells (Lucigen) according to the manufacturer’s protocol using a GenePulser (BioRad). To ensure no loss of representation, sufficient parallel transformations were performed using the same ligation reaction and plated onto 245 mm x 245 mm plates (Corning) with carbenicillin selection (50 ug/ml), which yielded 30-200 X library coverage. Colonies were scraped off plates and combined before plasmid DNA extraction using Endotoxin-Free Plasmid Maxiprep (Qiagen).

### Library sequencing and validation

To check library representation, synthesis fidelity, and bias, libraries were amplified and then deep sequenced. First, libraries were PCR amplified for 16 cycles using Phusion Flash High-Fidelity (Thermo) with primers to add adaptors for Illumina sequencing. For all libraries, PCR reactions were performed in duplicate and barcoded to allow quantification of bias introduced by PCR. Samples were sequenced on a MiSeq following the manufacturer’s protocol using a v3 150 cycle kit with 10 % PhiX (Illumina).

PCR replicates were demultiplexed using FASTX-Toolkit (Hannon Lab, CSHL) and adaptors were trimmed using cutadapt to leave only the sgRNA guide sequence^8^. Alignment of the guide sequence to the appropriate GeCKO library index was done using Bowtie with parameters to tolerate up to a single nucleotide mismatch^5^. The Bowtie alignment was then read into Matlab for further analysis (Mathworks). For all libraries, greater than 90 % of sgRNAs were represented with at least one sequencing read and the difference in representation between the 90^th^ and 10^th^ percentile sgRNAs was always less than 15-fold. Histograms of sgRNA distributions and cumulative sgRNA read counts for libraries in both lentiCRISPRv2 and lentiGuide-Puro are shown in Supplementary Figure 2 (human) and Supplementary Figure 3 (mouse).

### Vector maps, library sequences and reagent distribution

All lentiCRISPR plasmids and GeCKO libraries are available on Addgene. Please see http://genome-engineering.org/gecko/ for downloadable vector maps. See labeled sequences in Supplementary Data for more details.

The sgRNAs in the human and mouse v2 GeCKO libraries are available for download on the GeCKO website or from Addgene.

## Supplementary Data

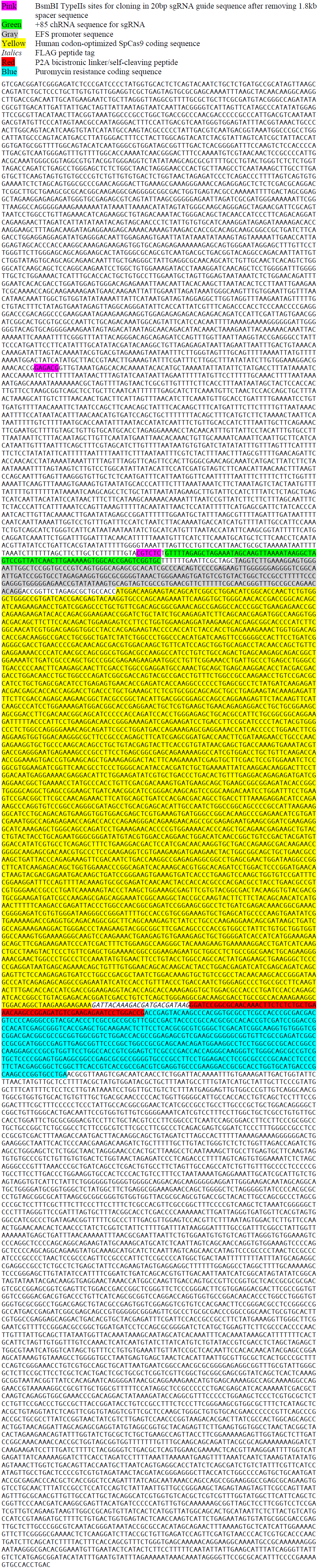
LentiCRISPR v2.

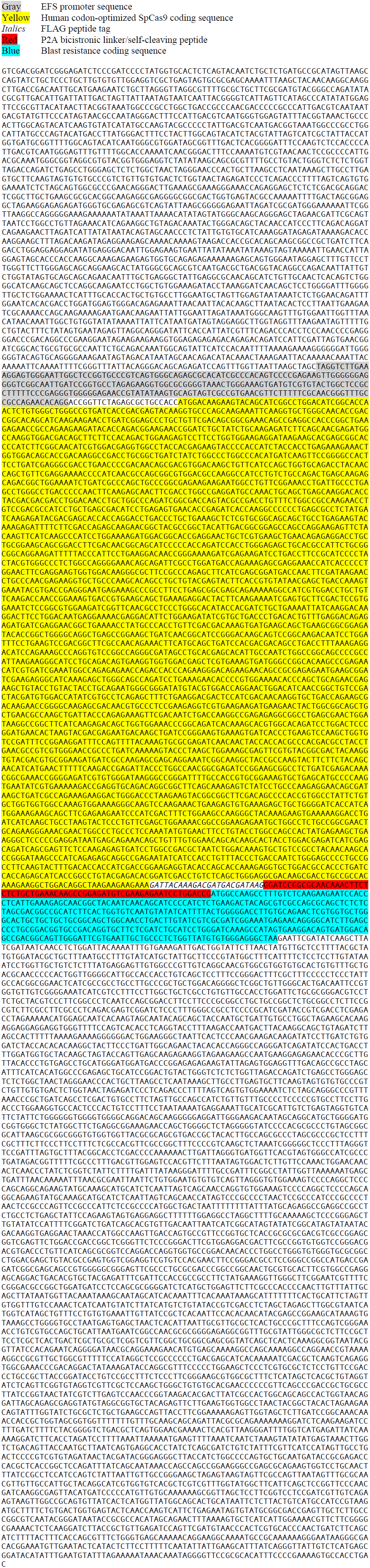
lentiCas9-Blast.

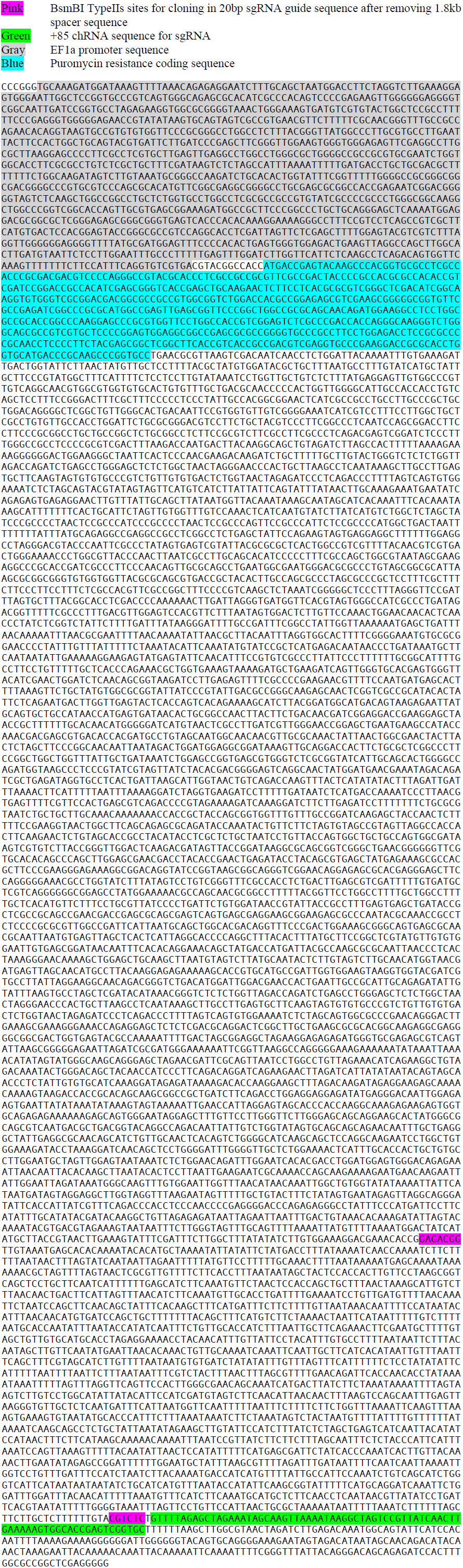
lentiGuide-Puro.

## References

1. Shalem, O. et al. Science 343, 84–87 (2014).

2. Wang, T., Wei, J. J., Sabatini, D. M. & Lander, E. S. Science 343, 80–84 (2014).

3. Koike-Yusa, H., Li, Y., Tan, E.-P., Velasco-Herrera, M. D. C. & Yusa, K. Nat Biotechnol 32, 267–273 (2013).

4. Zhou, Y. et al. Nature 509, 487–491 (2014).

5. Hsu, P. D. et al. Nat Biotechnol 31, 827–832 (2013).

6. Zhao, Y. et al. Sci Rep 4, 3943 (2014).

## References

1. Shalem, O. et al. Genome-scale CRISPR-Cas9 knockout screening in human cells. Science 343, 84–87 (2014).

2. Trapnell, C., Pachter, L. & Salzberg, S. L. TopHat: discovering splice junctions with RNA-Seq. Bioinformatics 25, 1105–1111 (2009).

3. Trapnell, C. et al. Differential gene and transcript expression analysis of RNA-seq experiments with TopHat and Cufflinks. Nat Protoc 7, 562–578 (2012).

4. Merkin, J., Russell, C., Chen, P. & Burge, C. B. Evolutionary dynamics of gene and isoform regulation in Mammalian tissues. Science 338, 1593–1599 (2012).

5. Langmead, B., Trapnell, C., Pop, M. & Salzberg, S. L. Ultrafast and memory-efficient alignment of short DNA sequences to the human genome. Genome Biol. 10, R25 (2009).

6. Hsu, P. D. et al. DNA targeting specificity of RNA-guided Cas9 nucleases. Nat Biotechnol 31, 827–832 (2013).

7. Kozomara, A. & Griffiths-Jones, S. miRBase: annotating high confidence microRNAs using deep sequencing data. Nucleic Acids Res 42, D68–73 (2014).

8. Martin, M. Cutadapt removes adapter sequences from high-throughput sequencing reads. EMBnet.journal 17, 10–12 (2011).

